# Physics-informed self-supervised learning enables spectra-free multiplexed imaging on standard fluorescence microscopes

**DOI:** 10.64898/2026.04.27.721244

**Authors:** Jiankai Xia, Jinhong Yan, Mingxuan Tang, Bin Zhao, Kun Chen

**Affiliations:** School of Optoelectronic Science and Engineering, University of Electronic Science and Technology of China, Chengdu 610054, China; Yangtze Delta Region Institute (Huzhou), University of Electronic Science and Technology of China, Huzhou 313000, China

## Abstract

Multiplexed fluorescence imaging is limited by spectral overlap and the small number of excitation or emission channels available on standard microscopes, restricting most laboratories to low-plex imaging. Here we introduce physics-informed spectra-free multiplexed imaging (PhySMI), a self-supervised framework for underdetermined spectral unmixing that enables highly multiplexed imaging without dense spectral measurements after training. By embedding the spectral forward-mixing process into a self-consistent architecture, PhySMI recovers physically plausible source decompositions from unlabeled data without paired ground-truth labels while suppressing stochastic acquisition noise. PhySMI resolves five subcellular structures from only three excitation channels, overcoming the conventional channel-number limit while preserving spectral fidelity and minimizing crosstalk (<0.5%). The framework also generalizes across imaging systems, enabling zero-shot deployment on standard fluorescence microscopes. In live cells, PhySMI enables fast five-color imaging of dynamic multi-organelle interactions with improved temporal resolution and reduced photobleaching and phototoxicity relative to conventional spectral imaging. These results establish a general strategy for physics-informed learning in underdetermined imaging inverse problems and represent a step toward a general-purpose framework for highly multiplexed fluorescence imaging on standard microscopy platforms.

## Introduction

Understanding how cellular organization gives rise to biological function requires imaging methods that can simultaneously resolve multiple molecular and organellar targets within the same cell^1, 2^. However, the multiplexing capability of conventional fluorescence microscopy remains fundamentally limited by the broad and overlapping spectra of fluorescent probes, restricting the number of distinguishable labels in a single experiment^3^. Fluorescence spectral imaging addresses this limitation by separating overlapping probes based on their spectral signatures, enabling the discrimination of four to six targets when combined with linear unmixing^4-6^. Most implementations rely on emission spectra, but acquiring spectral and spatial information simultaneously remains challenging in wide-field geometries^7, 8^. Consequently, emission-dispersed spectral imaging is typically performed using point scanning, limiting throughput and temporal resolution^9, 10^. Excitation spectral imaging provides an alternative by scanning excitation wavelengths across the full field of view while detecting fluorescence through a fixed emission channel, enabling faster multiplexed imaging with improved temporal resolution^11-14^.

Despite these advances, the widespread deployment of spectral imaging in biological laboratories remains limited by a fundamental constraint in spectral unmixing itself. Classical inverse methods require the problem to be determined or overdetermined, with at least as many spectral measurements as targets, and often more for robust decomposition^4, 5^. In addition, these solvers are highly susceptible to noise amplification, whereby stochastic fluctuations in low-photon regimes are propagated into the unmixed components, degrading spatial detail and structural fidelity^15-17^. Recent deep-learning-based unmixing methods, from supervised approaches trained on simulated ground truth to self-supervised emission-unmixing models, have improved reconstruction quality and robustness^18, 19^. However, they have not yet demonstrated experimental recovery of more fluorophore-labeled targets than acquired spectral channels, leaving the channel-number constraint unresolved for highly multiplexed live-cell imaging. Although this constraint can be partially relaxed when targets are spatially well separated^20^, such conditions are rarely satisfied in complex subcellular environments. As a result, highly multiplexed spectral imaging still depends on dense spectral acquisition, which in excitation spectral microscopy typically requires multiple discrete laser lines or broadband sources such as supercontinuum lasers^13, 14^. This increases system complexity and limits deployment to specialized platforms, whereas standard fluorescence microscopes in most laboratories typically provide only three to four excitation wavelengths.

Here we introduce physics-informed spectra-free multiplexed imaging (PhySMI), a self-supervised framework for underdetermined spectral unmixing and denoising that enables highly multiplexed imaging on standard fluorescence microscopes without dense spectral acquisition. PhySMI resolves up to five subcellular structures using only three excitation wavelengths, exceeding the conventional channel-number limit. By embedding the spectral forward-mixing process into self-supervised training, PhySMI constrains solutions to physically plausible manifolds and learns accurate source decompositions without paired ground-truth labels, while mitigating noise amplification and preserving structural detail. This physics-informed design also enables cross-system generalization, allowing zero-shot deployment across microscopes. By reducing the number of excitation measurements, PhySMI further enables faster multiplexed imaging while maintaining structural fidelity and low crosstalk. In live cells, this enables dynamic five-color imaging on a standard three-laser microscope with improved temporal resolution and reduced photobleaching and phototoxicity. Together, these advances broaden access to highly multiplexed live-cell fluorescence imaging in routine biological laboratories and establish a general strategy for physics-informed learning in underdetermined imaging inverse problems, representing a step toward a general-purpose framework for highly multiplexed fluorescence imaging on standard microscopy platforms.

## Results

### Reformulating spectral unmixing as a physics-informed forward inference problem

Fluorescence imaging is fundamentally limited by the extensive spectral overlap of fluorophores, as the broad profiles s_n_(*λ*) of different probes produce substantial crosstalk across channels, restricting the number of resolvable targets (Fig. 1a). Spectral imaging is governed by the forward mixing of latent biological structures into measured data. During acquisition (Fig. 1b–c), unknown object components *X* (*X*_1,_ … *X*_5_) are linearly combined to produce multispectral observations *Y*, following the linear mixing model (LMM): *Y* = **A***X + n*, where **A** denotes the spectral mixing matrix and *n* represents measurement noise. Each column of **A** encodes the spectral signature of a specific target, such that spatial information from individual structures is compressed into the observed spectral stack. Recovering these sources therefore constitutes an inverse problem.

**Figure 1.**
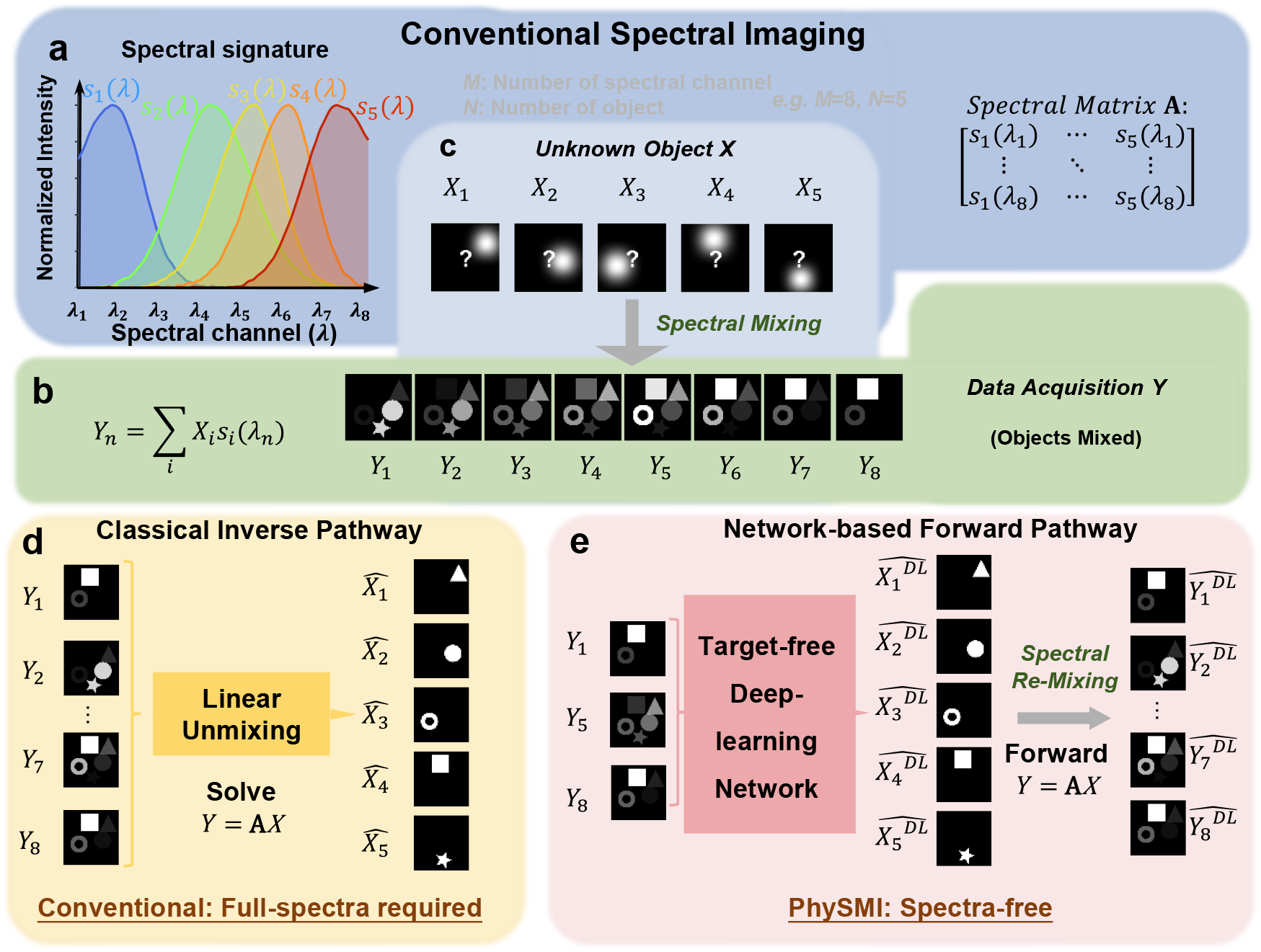
Principles of fluorescence spectral imaging and the PhySMI unmixing paradigm. (a) Spectral overlap in a congested fluorescence spectrum. Normalized emission profiles (s_1_(*λ*) - s_5_(*λ*)) exhibit substantial crosstalk, complicating the separation of multiple labeled targets. (b, c) Physical forward mixing and data acquisition. Latent biological structures (*X*_1_ *– X*_5_) are linearly combined during imaging according to the linear mixing model (LMM): *Y* = **A***X*, where *A* denotes the spectral mixing matrix. The underlying structures are unknown (indicated by “?”), and the measured data consist of mixed spectral observations (*Y*_1_ *– Y*_5_). (d) Conventional inverse pathway. Classical linear unmixing methods solve the inverse problem *Y* = **A***X* and require dense spectral sampling (*M* ≥ *N*) to recover the underlying sources, necessitating full-spectrum acquisition. (e) PhySMI forward inference paradigm. PhySMI infers *N* = 5 object sources from sparse spectral observations (*M* = 3 channels: *Y*_1_, *Y*_5_, *Y*_8_) without full spectral acquisition. A physics-informed re-mixing step enforces self-consistency by reconstructing the measured data via the forward model, establishing a closed-loop inference framework that bypasses the channel-number constraint of classical methods. The full network architecture is detailed in Fig. 2.

Conventional spectral unmixing addresses this problem using classical inverse solvers such as weighted least squares or non-negative least squares (NNLU) (Fig. 1d). These methods require the system to be determined or overdetermined (*M* ≥ *N*), and in practice rely on dense spectral sampling for stable decomposition, necessitating full-spectrum acquisition. This requirement increases hardware complexity and limits deployment on standard fluorescence microscopes. Here we instead reformulate spectral unmixing as a physics-informed forward inference problem that enables recovery in the underdetermined regime (*M* < *N*). Rather than directly inverting the mixing process, we embed the spectral forward model into a self-supervised framework (Fig. 1e), in which candidate source estimates are constrained by a differentiable re-mixing step that reconstructs the measured data. This enforces consistency with the physical mixing process and restricts solutions to a physically plausible subset, allowing recovery of high-dimensional source information from sparse observations without paired ground-truth labels. By removing the need for full-spectrum acquisition during inference, PhySMI enables spectra-free multiplexed imaging after training. In our implementation, PhySMI resolves *N* = 5 subcellular structures from only *M* = 3 excitation channels on standard fluorescence microscopes, establishing a practical framework for underdetermined spectral imaging in routine biological laboratories.

### A physics-informed self-supervised framework for underdetermined spectral unmixing

To realize the forward inference formulation described in Fig. 1, we developed a physics-informed self-supervised framework for spectral unmixing in the underdetermined regime (*M* < *N*). In this setting, multispectral measurements arise from the linear mixing of multiple fluorophores into a compressed spectral space, resulting in an ill-posed inverse problem with infinitely many feasible solutions. The absence of paired ground-truth labels in live-cell imaging further precludes conventional supervised learning, necessitating an alternative strategy that constrains solutions through intrinsic physical and structural priors.

We address this challenge using a dual-branch self-consistent training paradigm (Fig. 2a). During training, two independently sampled spectral subsets (*Y*_*branch α*_, *Y*_*branch β*_) are processed by a weight-sharing unsupervised unmixer. Instead of relying on external labels, the model enforces consistency through cross-branch reconstruction: source estimates inferred from one branch are required to explain the measurements of the other after passing through the corresponding spectral forward model. This design exploits the statistical independence of acquisition noise across branches while preserving the shared biological signal, driving the network to learn noise-invariant representations of the underlying structures. At inference, a single branch is sufficient for source recovery. Model convergence is governed by three complementary constraints that restrict solutions to physically and biologically plausible manifolds. A reconstruction loss enforces spectral consistency by requiring that predicted sources, when re-mixed through the fixed spectral mixing matrix, reproduce the measured observations. An adversarial loss incorporates morphological priors through an unpaired biological structure library, suppressing structurally implausible solutions. A cross-branch similarity loss enforces cyclic consistency between branches, promoting agreement on deterministic signals while attenuating stochastic noise^21, 22^. Together, these constraints replace the need for paired supervision and stabilize learning in the rank-deficient regime.

**Figure 2.**
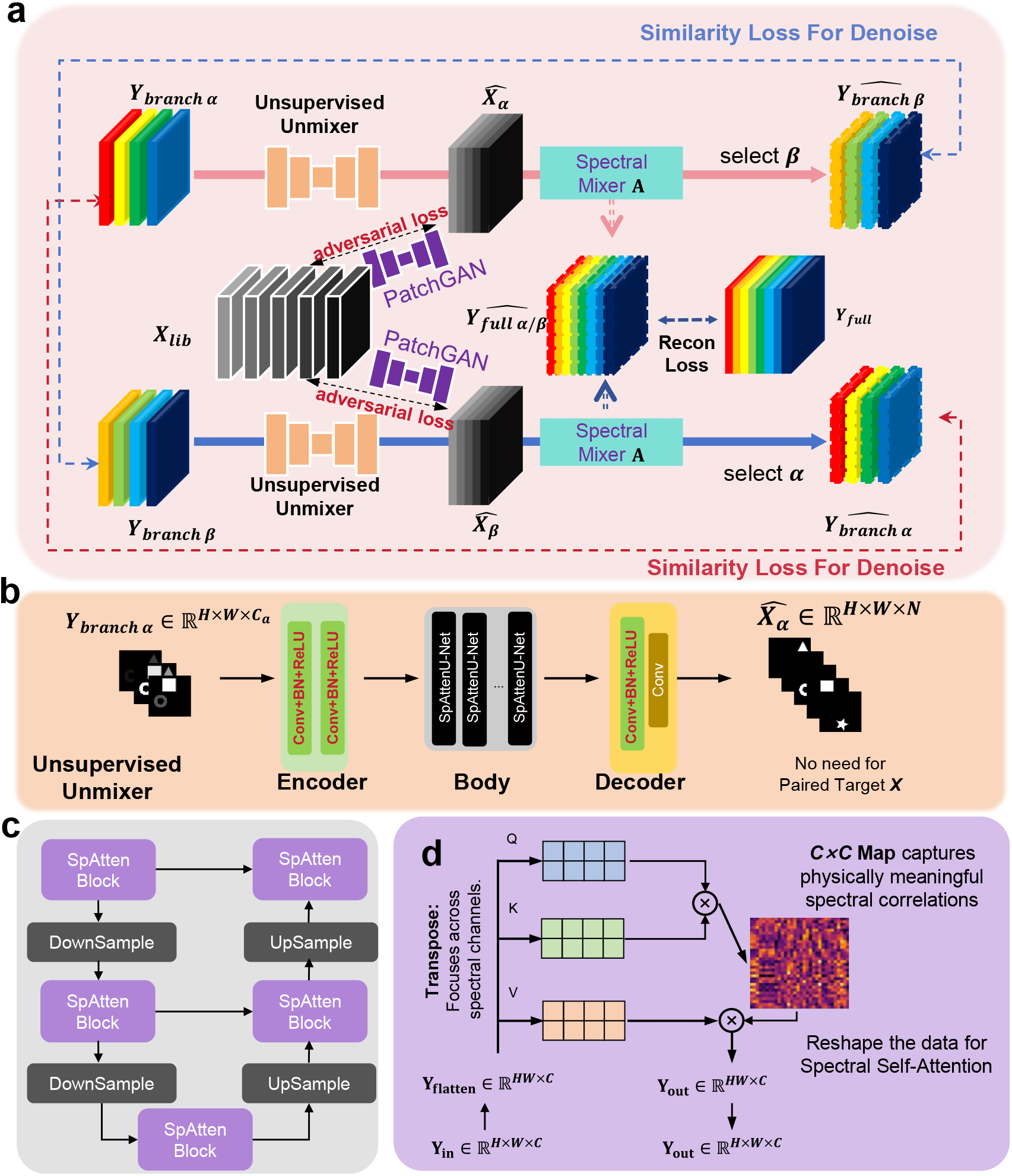
PhySMI Framework Architecture. (a) Dual-branch self-consistent training paradigm. A weight-sharing unsupervised unmixer ℱ_θ_ processes two independently sampled sparse inputs (*Y*_*branch α*_, *Y*_*branch β*_). during training. Training is guided by three complementary constraints: (i) reconstruction loss (ℒ_*rec*_), which enforces physical consistency by re-mixing the inferred sources through the fixed spectral mixing matrix **A**; (ii) adversarial loss (ℒ_*adv*_), which incorporates morphology priors using PatchGAN discriminators and an unpaired biological structure library *X*_*lib*_; and (iii) cross-branch similarity loss (ℒ_*sim*_), which promotes noise-invariant representations via cyclic reconstruction between the two branches. (b) Unsupervised unmixer. An encoder–decoder network that reconstructs *N* source components from a single sparse input without paired ground-truth labels. (c) Hierarchical SpAtten U-Net backbone. A multi-scale U-Net architecture augmented with spectral self-attention (SpAtten) blocks at each resolution level to capture long-range dependencies across spectral channels. (d) Spectral self-attention mechanism. Input features **Y**_**in**_ are reshaped into channel-wise tokens to construct a *C*×*C* spectral correlation map, enabling the network to model physically meaningful spectral relationships and guide nonlinear source separation.

The unsupervised unmixer is implemented as a hierarchical encoder–decoder network with a multi-scale U-Net backbone (Fig. 2b–c). To disentangle spectrally overlapping signals, we introduce spectral self-attention (SpAtten) blocks that operate along the channel dimension (Fig. 2d)^23^. By constructing channel-wise correlation maps, this mechanism captures physically meaningful spectral relationships and enables adaptive recombination of undersampled spectral features, facilitating accurate source separation under severe spectral overlap. Together, these components establish a framework for underdetermined spectral unmixing and self-supervised denoising, enabling stable recovery of multiplexed fluorescence signals from sparse spectral measurements without paired ground-truth labels.

### Accurate and interpretable recovery with PhySMI under spectral undersampling

To quantitatively evaluate PhySMI in the underdetermined regime, we constructed a simulated spectral imaging dataset with known ground truth (Fig. 3a). Five singly labeled subcellular structures in fixed COS-7 cells—mitochondria, nucleus, lipid droplets (LDs), microtubules, and actin—were acquired as source components. Synthetic spectral images were generated from these structures (*X*) and the measured excitation spectra of the corresponding fluorophores (**A**) using the linear mixing model *Y* = **A***X* + *n*, with additive Gaussian noise (5% of signal amplitude) to emulate stochastic acquisition noise. From the resulting full spectral stack (Y_*full*_, 8 channels), a sparse three-channel subset was selected as input to PhySMI (Fig. 3b), mimicking standard fluorescence microscope configurations. Despite severe spectral undersampling (*M* = 3, *N* = 5), PhySMI accurately recovered all five subcellular structures (Fig. 3c), demonstrating its ability to resolve multiplexed signals beyond the conventional channel-number limit.

**Figure 3.**
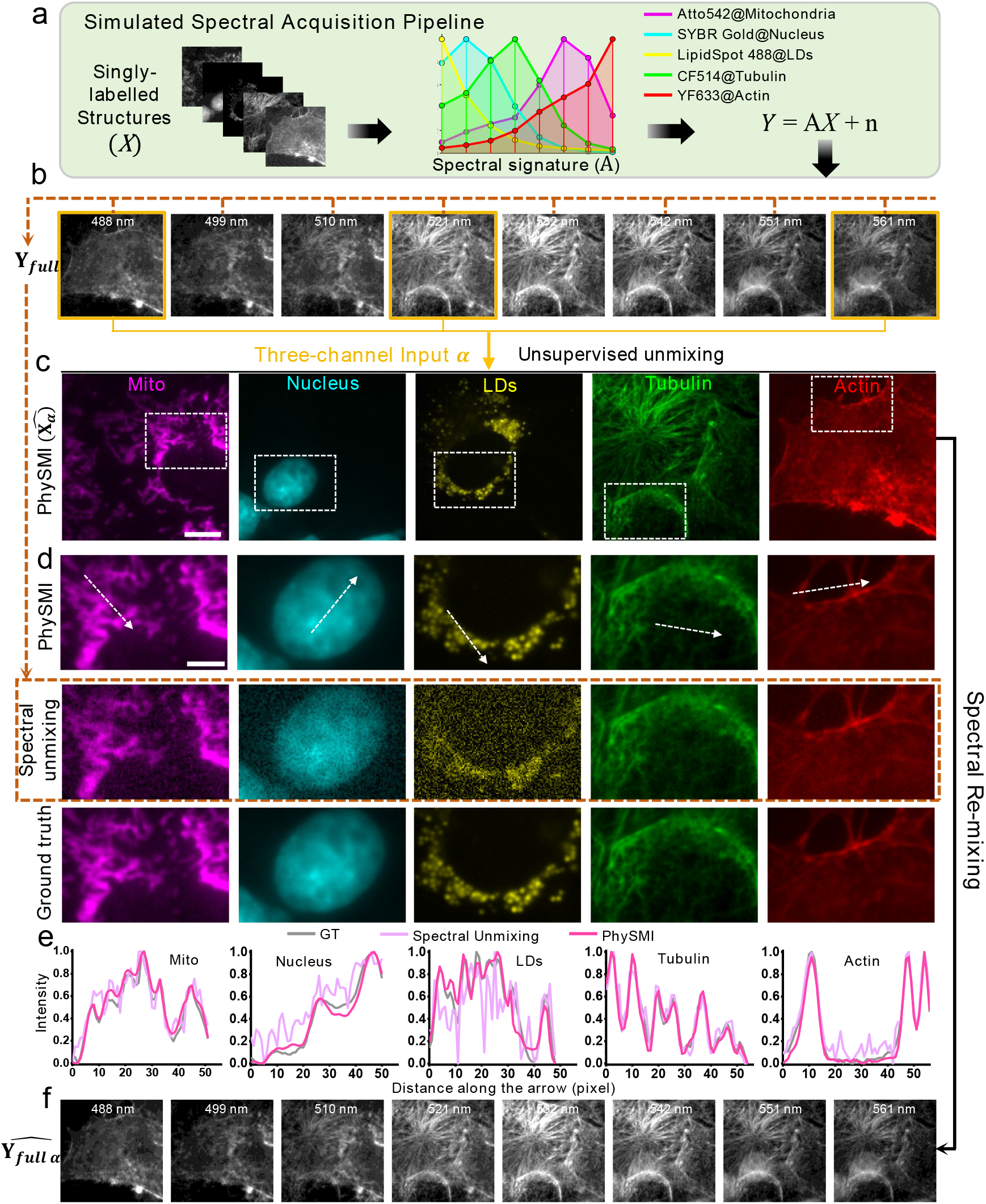
Validation of the PhySMI framework using simulated spectral data. (a) Simulated spectral acquisition pipeline. Five singly labeled subcellular structures in fixed COS-7 cells—mitochondria (ATTO542), nucleus (SYBR Gold), lipid droplets (LipidSpot 488), microtubules (CF514), and actin (YF633)—were used as source components. Synthetic multispectral data were generated by combining these images with their excitation spectral signatures using the linear mixing model *Y*=**A***X*+n, with additive Gaussian noise (5% of signal amplitude) applied at each wavelength. (b) Full-spectrum observations and sparse input. An 8-channel spectral stack (Y_*full*_, 488, 499, 510, 521, 532, 542, 551, 561nm) was generated, from which a sparse three-channel subset (*Y*_*branch α*_) was selected as input to PhySMI. (c) PhySMI reconstruction. Unmixed results 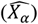 of the five subcellular structures recovered from the three-channel input. (d) Comparison with linear unmixing and ground truth. Representative regions (with dashed boxes in c) show that spectral unmixing suffers from noise amplification and crosstalks, whereas PhySMI recovers structures with high fidelity, closely matching the ground truth. (e) Line profile analysis. Normalized intensity profiles along the dashed lines in (d) highlight improved signal separation and reduced crosstalk with PhySMI. (f) Physical self-consistency. Re-mixing the inferred components using the spectral matrix **A** yields a reconstructed full-spectrum stack 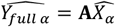 that closely matches the original measurements, confirming preservation of the underlying spectral physics. Scale bar: 10 μm (b, c, f) and 5 μm (d)

We next compared PhySMI with conventional linear unmixing under identical conditions (Fig. 3d). Although linear unmixing recovered some structures reasonably well, it exhibited pronounced noise amplification for lower-SNR components, particularly the nucleus and LDs, resulting in reduced structural fidelity and signal leakage. By contrast, PhySMI reconstructed all five structures with high fidelity, preserving fine structural details while effectively suppressing noise amplification. Line profiles extracted from representative regions (Fig. 3e) further confirm improved signal separation and reduced crosstalk. To verify physical consistency, the recovered structures were re-mixed using the spectral mixing matrix **A**. The reconstructed full-spectra stack closely matched the original observations across all spectral channels (Fig. 3f), indicating that PhySMI preserves the underlying spectral physics while operating in the underdetermined regime.

To determine how the key constraints in PhySMI contribute to recovery performance, we performed a systematic ablation study using the same simulated dataset (Table 1). The full model achieved the best overall performance (PSNR = 39.97 dB; SSIM = 0.977) and the lowest spectral angle mapper (SAM), indicating high spectral fidelity and strong physical self-consistency. These values also exceeded those obtained with conventional eight-channel NNLU (PSNR = 33.82 dB; SSIM = 0.861), providing quantitative support for the improved reconstruction fidelity observed in Fig. 3d. Removing the physical reconstruction constraint (ℒ_*rec*_) caused the most severe collapse (PSNR = 20.76 dB; SAM = 25.05), demonstrating that the spectral mixing model serves as an essential physical anchor for valid source recovery. The morphological prior (ℒ_*adv*_) improved structural fidelity by suppressing implausible reconstructions, whereas the cross-branch similarity constraint (ℒ_*im*_) further refined recovery by attenuating stochastic acquisition noise and enforcing consistency across independent spectral observations. Together, these results show that physical, structural, and statistical constraints are jointly required for stable recovery in the underdetermined regime.

**Table 1.** Quantitative Ablation Study of PhySMI Framework.

| Model Case | Physical Recon<br>($\mathcal{L}_{rec}$) | Morphological Prior ($\mathcal{L}_{adv}$) | Dual-Branch Consistency<br>($\mathcal{L}_{sim}$) | PSNR (dB)<br>↑ | SSIM ↑ | SAM ↓ |
| --- | --- | --- | --- | --- | --- | --- |
| I (Base) | ✓ | / | / | 36.32 | 0.907 | 4.28 |
| II | ✓ | / | ✓ | 36.56 | 0.938 | 3.88 |
| III | ✓ | ✓ | / | 39.02 | 0.971 | 2.93 |
| Full (Ours) | ✓ | ✓ | ✓ | **39.97** | **0.977** | **2.61** |
| IV (No Phys.) | / | ✓ | / | 20.76 | 0.664 | 25.05 |
| NNLU (8 ch.) | / | / | / | 33.82 | 0.861 | / |

To further understand how PhySMI resolves multiple targets from sparse inputs, we performed a perturbation-based attribution analysis (Supplementary Fig. S1). By systematically masking individual spectral channels and quantifying the resulting loss of reconstruction fidelity, we derived channel importance maps that reveal the model’s internal inference strategy. Each input channel not only supports specific targets but also constrains others to suppress crosstalk. For example, a channel that strongly supports actin reconstruction also serves as a critical disambiguation cue for spectrally overlapping structures such as microtubules; masking this channel leads to pronounced signal leakage between components, indicating that accurate unmixing depends on the synergistic integration of all available spectral cues.

Consistent with this mechanism, PhySMI also exhibits enhanced robustness to stochastic acquisition noise compared with conventional spectral unmixing. Controlled simulations under increasing noise levels show that conventional unmixing suffers from strong noise amplification and rapidly increasing crosstalk, whereas PhySMI maintains stable reconstruction with minimal degradation (Supplementary Fig. S2–S3). These findings indicate that PhySMI does not rely on simple intensity mapping, but instead learns a physically grounded representation of spectral correlations. By capturing invariant spectral relationships across channels, the model reconstructs high-dimensional source information from undersampled observations while suppressing noise-driven degradation, providing a mechanistic basis for its ability to generalize beyond the spectral configurations used during training. This behavior is consistent with the physics-informed forward inference formulation introduced in Fig. 1.

### Zero-shot deployment of PhySMI on standard fluorescence microscopes

To evaluate whether PhySMI generalizes across imaging systems, we tested its ability to resolve five subcellular structures in fixed cells on a standard three-laser fluorescence microscope using a model trained exclusively on a supercontinuum-based excitation spectral imaging system (Fig. 4). We first assessed performance on the excitation spectral imaging system by comparing PhySMI using a sparse three-channel input (488, 532, 561 nm) with conventional spectral unmixing based on the full eight-channel dataset (Fig. 4a and Supplementary Fig. S4, see Methods). Although linear unmixing is provided with complete spectral information, it exhibits increased noise and reduced structural definition in lower-SNR channels, particularly for the nucleus and LDs. By contrast, PhySMI reconstructs all five structures with high fidelity from only three excitation channels. Zoomed-in views highlight effective suppression of noise and artifacts, while line profiles across LDs show that PhySMI preserves fine structural features that are blurred in conventional unmixing. Applying the same five-fluorophore PhySMI unmixing procedure to samples singly labeled with each fluorophore showed an average target-to-target crosstalk below 0.5% (Supplementary Fig. S5), further supporting accurate structure separation under sparse three-channel excitation.

**Figure 4.**
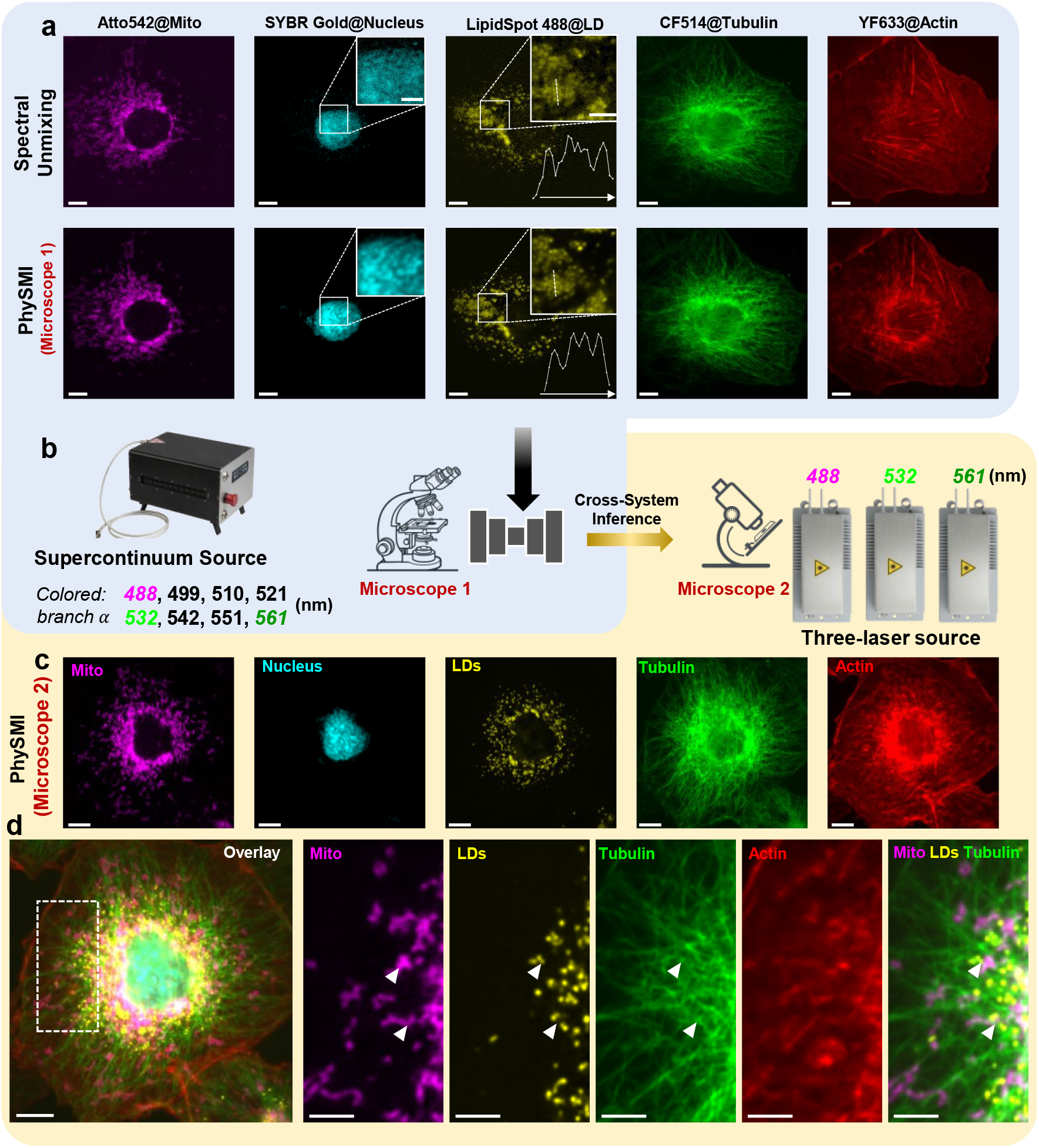
Zero-shot deployment of PhySMI on standard fluorescence microscopes. (a) Experimental comparison between full-spectrum spectral unmixing and sparse-input PhySMI. Five subcellular structures in fixed COS-7 cells—mitochondria (ATTO542), nucleus (SYBR Gold), LDs (LipidSpot 488), microtubules (CF514), and actin (YF633)—were imaged on System 1 using full 8-channel excitation spectral acquisition (488, 499, 510, 521, 532, 542, 551, and 561 nm). Conventional spectral unmixing using the full spectral stack is compared with PhySMI applied to a sparse three-channel subset (488, 532, 561 nm). Despite using fewer measurements (*M* = 3), PhySMI reconstructs all five structures with improved SNR and structural fidelity. Zoomed-in insets highlight reduced noise and artifact suppression in the nucleus and LD channels. Line profiles in the LD inset further illustrate enhanced spatial resolution and reduced blurring in the PhySMI reconstruction. (b) Cross-system transfer scheme. Schematic of model transfer from a supercontinuum-based excitation spectral imaging system (Microscope 1) to a standard three-laser fluorescence microscope (Microscope 2; 488, 532, 561 nm). The model is trained on Microscope 1 using spectral subsampling aligned to the excitation wavelengths available on Microscope 2. (c) Zero-shot deployment on a standard fluorescence microscope. Representative five-color image of a fixed COS-7 cell acquired on Microscope 2 and resolved using a model trained exclusively on Microscope 1. Despite the hardware shift and the underdetermined condition (*M* < *N*), PhySMI recovers all five subcellular structures with minimal target-to-target crosstalk. (d) Merged image and spatial organization. Overlay and zoomed-in views reveal spatial colocalization and contact relationships among mitochondria, LDs, actin, and microtubules. Arrowheads indicate representative contact sites. Scale bars: 10 μm (a, c, d) and 5 µm (zoomed regions in a, d).

We next evaluated zero-shot transfer to a standard fluorescence microscope. As illustrated in Fig. 4b, the model was trained using spectral subsampling aligned to the three excitation wavelengths available on the target system and then directly applied, without retraining, to data acquired on the standard microscope (Supplementary Fig. S6, see Methods). Despite the hardware shift and the underdetermined condition (*M* = 3, *N* = 5), PhySMI successfully resolves all five subcellular structures with minimal crosstalk (Fig. 4c). The merged images further reveal coherent spatial organization and interactions among these structures (Fig. 4d), including representative contact sites between mitochondria, LDs, and microtubules.

This cross-system robustness arises from the physics-informed design of PhySMI, which explicitly leverages the physical basis of excitation spectral imaging. The excitation spectral mixing matrix **A** is governed primarily by the intrinsic absorption and radiative responses of fluorophores and is therefore largely insensitive to detector sensitivity, emission filters, and other system-specific detection characteristics. As long as excitation wavelengths and relative illumination conditions are matched, the spectral mixing relationships remain consistent across microscopes. By learning these invariant excitation-dependent responses, PhySMI enables zero-shot deployment of highly multiplexed imaging on standard fluorescence microscopes without specialized spectral acquisition hardware.

### PhySMI enables fast five-color live-cell imaging on standard fluorescence microscopes

We next evaluated whether PhySMI enables fast multiplexed imaging of dynamic processes in live cells using standard fluorescence microscopes. Using a three-laser system, we applied PhySMI to track multi-organelle dynamics from only three excitation channels (Fig. 5a, Supplementary Fig. S7, Supplementary Video 1 and Video 2). The framework resolves five subcellular structures—including mitochondria, LDs, peroxisomes, nucleus, and lysosomes—providing a comprehensive view of the subcellular landscape (Fig. 5b). Reducing the number of excitation measurements directly improves imaging speed. Compared with conventional excitation spectral imaging requiring dense multi-channel acquisition (e.g., eight channels), the three-channel implementation increases the effective temporal resolution by ∼2.7-fold at a fixed camera rate. At 10 Hz acquisition, this corresponds to 0.3 s per five-color image, enabling the capture of transient multi-organelle interactions that would be undersampled under full spectral acquisition.

**Figure 5.**
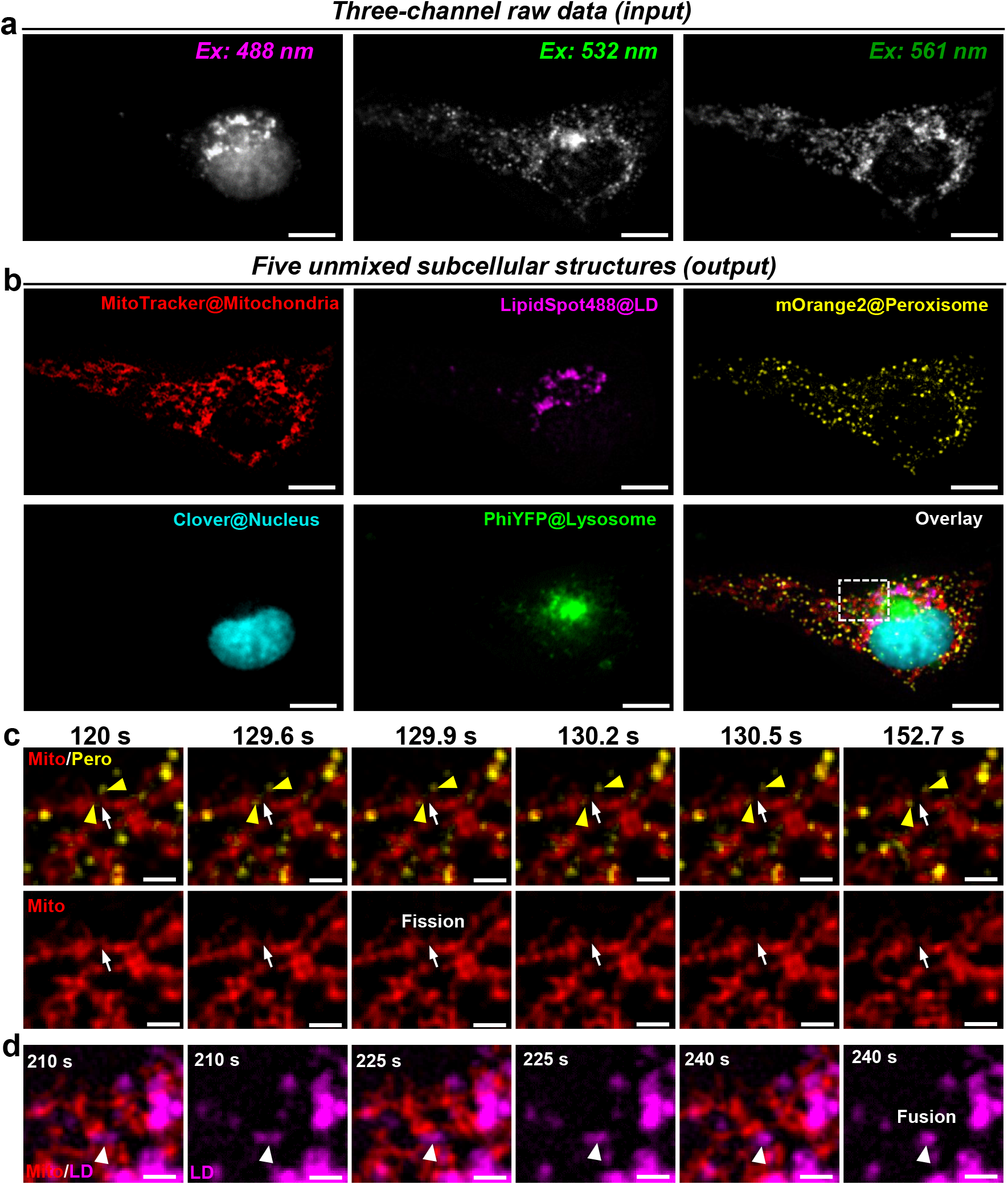
PhySMI enables fast five-color live-cell imaging of dynamic subcellular interactions on a standard fluorescence microscope. (a) Three-channel raw inputs. Live-cell fluorescence images acquired at 488, 532, and 561 nm excitation on a standard three-laser fluorescence microscope. These three channels serve as sparse input measurements for PhySMI. (b) PhySMI-resolved five-color imaging. Reconstruction of five subcellular structures in live cells, including mitochondria (MitoTracker), LDs (LipidSpot 488), peroxisomes (mOrange2), nucleus (Clover), and lysosomes (PhiYFP), together with the merged overlay. (c) Mitochondrial fission event. Magnified time-lapse views of the boxed region in (b) show a mitochondrial fission event (white arrows) occurring at a site associated with peroxisome contact (yellow arrowheads), illustrating multi-organelle coordination captured by PhySMI. (d) Dynamic organelle interactions. Time-lapse imaging of mitochondria–LD contacts and a LD fusion event (arrowheads), demonstrating simultaneous visualization of dynamic processes across multiple organelles. All live-cell imaging was performed at a camera acquisition rate of 10 Hz, corresponding to a temporal resolution of 0.3 s per five-color PhySMI image. Scale bars: 10 μm (a, b) and 2 µm (c, d).

Time-lapse imaging reveals dynamic processes, including mitochondrial fission at sites of peroxisome contact (Fig. 5c)^24, 25^ and coordinated LD fusion and interactions with the mitochondrial network (Fig. 5d)^26^. These results demonstrate that PhySMI preserves both structural detail and temporal fidelity across channels. In addition to improving temporal resolution, reducing excitation cycles lowers the cumulative light dose, mitigating photobleaching and phototoxicity. By resolving *N* = 5 targets from *M* = 3 measurements, PhySMI shifts multiplexing from hardware-intensive acquisition to physics-informed computational inference, enabling efficient, high-dimensional imaging of dynamic subcellular interactions on standard fluorescence microscopes.

## Discussion

In this work, we present PhySMI, a physics-informed self-supervised framework that enables multiplexed fluorescence imaging beyond the conventional channel-number limit. By reformulating spectral unmixing as a physics-informed forward inference problem, PhySMI recovers multiple fluorophore-labeled targets from undersampled excitation measurements while preserving spectral fidelity and suppressing crosstalk. This capability is achieved without paired ground-truth labels and extends across imaging systems, enabling zero-shot deployment on standard fluorescence microscopes. Together, these results establish a practical route toward high-plex imaging that is no longer coupled to dense spectral acquisition or specialized hardware.

A central feature of PhySMI is that its robustness and transferability arise from the same physics-informed design. In classical spectral unmixing, measurement noise is propagated through linear inversion, often leading to noise amplification and loss of structural fidelity, particularly in low-photon regimes. By contrast, PhySMI constrains recovery through forward-model consistency and cross-branch self-supervision, restricting solutions to physically plausible manifolds while attenuating stochastic noise. The same formulation also explains its cross-system generalization. In excitation spectral imaging, the mixing process is governed primarily by the intrinsic absorption and radiative responses of fluorophores across excitation wavelengths and is therefore largely independent of the detection pathway. By learning these invariant excitation-dependent responses, PhySMI anchors inference to physical properties that are preserved across microscopes, enabling robust zero-shot transfer. In contrast, emission-based spectral imaging depends more strongly on the detection pathway, making such transfer inherently more challenging. Taken together, these results illustrate how embedding physical constraints into learning-based models improves both robustness and interpretability in ill-posed imaging inverse problems.

Recent deep-learning approaches have demonstrated high-dimensional *mn imlmco* visualization of intracellular structures from a universal membrane stain by learning morphology- and ratio-based organelle segmentation^27^. Although conceptually powerful, these methods address organelle classification and segmentation rather than spectral unmixing of simultaneously mixed fluorophore signals, and therefore tackle a problem distinct from the reduced-channel multiplexing framework presented here. They also rely on morphology-informed predictions and structure-specific training strategies. By contrast, PhySMI directly recovers fluorophore-resolved images under a physical spectral mixing model, enabling reduced-channel multiplexed imaging of specifically labeled targets without organelle-specific segmentation priors or task-specific supervision. These approaches are best viewed as complementary: morphology-driven inference can expand apparent labeling capacity from a single dye, whereas physics-informed unmixing enables simultaneous recovery of spectrally encoded targets under mixed-signal conditions.

Beyond its methodological advances, PhySMI provides direct biological utility by enabling five-color live-cell imaging of dynamic multi-organelle interactions on standard fluorescence microscopes. In our demonstrations, this includes mitochondrial fission associated with peroxisome contact, as well as LD fusion and interactions with the mitochondrial network. By reducing the number of required excitation measurements, PhySMI improves temporal resolution while lowering cumulative light exposure, thereby mitigating photobleaching and phototoxicity in live-cell imaging. The current framework assumes a known or calibrated spectral mixing model; to facilitate practical adoption, we provide a curated GitHub resource that includes excitation spectra for commonly used fluorophores together with the deep-learning models used in this work. Performance may degrade when spectral signatures are highly degenerate or when signal levels are extremely low. Future work could extend this framework through adaptive spectral calibration, probe design optimization, or hybrid strategies that combine physics-informed unmixing with data-driven priors. Overall, these results demonstrate that physics-informed learning provides a general and scalable strategy for overcoming fundamental constraints in multiplexed fluorescence imaging, representing a step toward a general-purpose framework for highly multiplexed imaging on standard microscopy platforms.

## Methods

### Optics setup

#### Excitation spectral imaging system with a supercontinuum laser source

A custom excitation spectral imaging system was used to acquire reference datasets for PhySMI training and validation^28^. Broadband excitation was provided by a supercontinuum laser (SC-Pro-M-40, YSL Photonics). The beam was linearly polarized using a polarizing beam splitter (PBS9012, Union Optic) and a linear polarizer (SHP1025, Union Optic), filtered by a short-pass filter (FESH0750, Thorlabs), and directed into an acousto-optic tunable filter (AOTF; 97-03151-01, Gooch & Housego). The selected excitation beam was expanded, collimated, and coupled into an inverted fluorescence microscope (Nikon Ti2-U), where it was focused onto the back focal plane of an air objective (CFI Plan Fluor, 60×, NA 0.85). An internal 1.5× magnification module was used, resulting in an effective system magnification of 90×. The excitation filter, dichroic mirror, and emission filter were FF01-505/119 (Semrock), FF573-Di01 (Semrock), and ET570lp (Chroma Technology), respectively. Fluorescence images were recorded using an EMCCD camera (iXon Ultra 897, Andor) at 512 × 512 pixels and 10 frames per second (fps), corresponding to an approximately 91 × 91 μm^2^ field of view.

For excitation spectral acquisition, an 8-channel RF synthesizer (97-03926-14, Gooch & Housego) drove the AOTF to sequentially select eight excitation wavelengths: 488, 499, 510, 521, 532, 542, 551, and 561 nm. Wavelengths were switched on a frame-by-frame basis, producing one complete excitation spectral stack every eight frames. A multifunction I/O board (PCI-6733, National Instruments) synchronized camera exposure with AOTF switching. For cellular imaging, the excitation power was maintained at approximately 25 μW per wavelength at 10 fps.

#### Standard three-laser fluorescence microscope for cross-system validation

Three-channel fluorescence imaging was performed on a Nikon Ti2-E inverted fluorescence microscope for cross-system validation^29^. The system was equipped with three excitation lasers: 488 nm (OBIS 488 LX, Coherent), 532 nm (NPLH-532-OEM, PL Optics), and 561 nm (MGL-FN-561, CNI Laser). The beams were collinearly combined and directed into the microscope through a dichroic mirror (ZT561rdc, Chroma), then focused onto the back focal plane of the same air objective (CFI Plan Fluor, 60×, NA 0.85). An internal 1.5× magnification module yielded an effective magnification of 90× at the camera plane.

For acquisition, excitation wavelengths were sequentially switched at 488, 532, and 561 nm using another AOTF (97-03151-01, Gooch & Housego), producing one complete excitation cycle every three frames. Fluorescence emission was separated from excitation by a long-pass filter (ET570lp, Chroma) and recorded using an sCMOS camera (Zyla 4.2, Andor). Images were acquired at 512 × 512 pixels after 2 × 2 binning at 10 fps, corresponding to an approximately 74 × 74 μm^2^ field of view. A multifunction I/O board (PCI-6733, National Instruments) synchronized camera exposure with excitation switching. For cellular imaging, the excitation power was maintained at approximately 25 μW per wavelength.

### Network architecture and implementation

#### Dual-branch self-consistent training paradigm

To address the non-uniqueness of spectral unmixing in the rank-deficient regime (*M* < *N*), we implemented a dual-branch self-consistent training paradigm (Fig. 2a). During training, two sparse spectral subsets (*Y*_*branch α*_, *Y*_*branch β*_) were processed by a weight-sharing Unsupervised Unmixer. The dual-branch design establishes a mutual supervision loop that helps decouple stochastic acquisition noise from deterministic biological signals. Each branch independently predicts the latent components 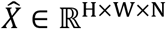, allowing single-branch inference during deployment.

Cross-branch consistency was enforced through a reconstruction mechanism (Fig. 2a, dashed feedback lines). Specifically, the unmixed estimate from one branch (for example, 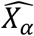) was required to reconstruct the raw observations of the other branch (*Y*_*branch β*_) after passing through the corresponding spectral selection operator (**A**_*β*,_ sub-matrices of **A** corresponding to the spectral channels selected for *β* branch). Because acquisition noise is statistically independent across branches, this design encourages the network to suppress branch-specific fluctuations and retain only shared biological signals.

#### Hierarchical SpAttenU-Net backbone

The Unsupervised Unmixer was implemented as a symmetric encoder–decoder network, termed Hierarchical SpAttenU-Net (Fig. 2b-c). The encoder transforms spectral–spatial inputs into a latent feature representation, and the decoder reconstructs the unmixed source components. A multi-scale hierarchical structure was used to capture diverse subcellular morphologies, ranging from filamentous microtubules to volumetric nuclei. Within this backbone, standard convolutional blocks were replaced by Spectral Self-Attention (SpAtten) modules to facilitate non-linear disentanglement of overlapping spectral signals.

#### Spectral Self-Attention (SpAtten) mechanism

The SpAtten module explicitly models spectral recombination (Fig. 2d). Inspired by the MST++^23^ framework, attention was applied along the spectral channel dimension rather than the spatial domain. Input feature maps *Y*_*in*_ ∈ ℝ^*H*×*W*×*C*^ were reshaped into channel-wise tokens to compute a *C* × *C* spectral correlation map using Query (*Q*), Key (*K*), and Value (*V*) matrices:

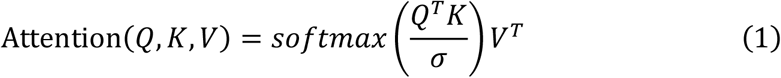

where σ is a learnable scaling factor. This correlation map captures relationships among spectral bands and was used to adaptively re-weight spectral features during source reconstruction.

#### Physical Reconstruction Loss

To ensure prediction fidelity to the linear mixing physics, the framework incorporates the fixed physical mixing matrix A as a differentiable layer within the computational graph. This ensures that the physical constraints defined by the LMM are directly coupled with the network’s optimization, allowing the gradients of the reconstruction error to be backpropagated through the physical model to refine the unmixing weights of the backbone. the predicted source 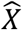 is re-mixed to reconstruct the full multispectral observation *Y*_*full*_:

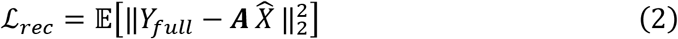

#### Adversarial Morphological Prior Loss

In the absence of paired ground-truth (GT), we utilize an unpaired morphological library (*X*_*lmb*_) via PatchGAN discriminators. As indicated in Fig. 2a, *N* independent discriminators are deployed to distinguish between the predicted component 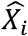 and the corresponding biological template from *X*_*lmb,m*_:

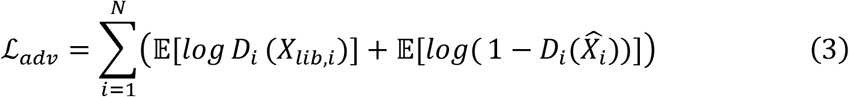

#### Similarity Loss for Cross-branch Reconstruction and denoising

The core of our self-supervised stability lies in the Cross-branch Reconstruction mechanism. Unlike conventional similarity constraints that operate in latent space, ℒ_*sim*_ enforces physical consistency by requiring the unmixed components from one branch to explain the raw spectral observations of the parallel branch.Specifically, the unmixed estimate 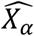 derived from the input *Y*_*branch α*_ is re-mixed using the sub-matrix **A**_*β*_ to generate a prediction of the *β*-branch observations, denoted as 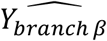. The loss is formulated as:

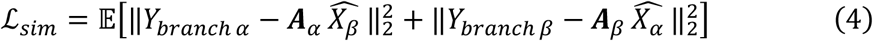

where the reconstructed branch inputs are defined as:

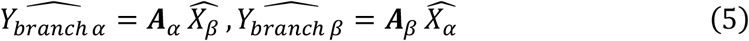

Here, **A**_*α*_ and **A**_*β*_ represent the fixed physical sub-matrices of **A** corresponding to the spectral channels selected for each branch.

#### Training and optimization

Training was performed using the dual-branch architecture with spectral subsets configured to match standard discrete laser lines: 488, 532, and 561 nm for branch *α* and 521, 542, 551 nm for branch *β*. The model was trained on 128×128 spatial patches with a batch size of 4 for 200 epochs. The total loss function was defined as a weighted sum of the reconstruction, similarity, and adversarial losses, with weights set to *λ*_rec_ = 2000.0, *λ*_sim_ = 1000.0 (denoising), *λ*_adv_ = 1.0. Optimization was performed using the Adam optimizer with (β_1_=0.9, β_2_=0.999). The initial learning rate for the generator was 2×10^−4^ and decayed exponentially to 4×10^−6^. The spectral endmember matrix was fixed throughout training, while the abundance estimation network was optimized. Input data were normalized using a fixed scaling factor before network input.

### Cell culture and sample preparation

#### Cell culture and fixation

COS-7 cells (Cell Resource Center, IBMS, CAMS/PUMS) were cultured on 18-mm glass coverslips in 12-well plates in Dulbecco’s Modified Eagle Medium (DMEM; Gibco) supplemented with 10% fetal bovine serum, and 1× non-essential amino acids at 37 °C in 5% CO_2_. After 24 h, cells were fixed with 3% paraformaldehyde and 0.1% glutaraldehyde in phosphate-buffered saline (PBS), followed by two washes with 0.1% sodium borohydride in PBS to quench residual aldehydes, and three washes with PBS (10 min each).

#### Plasmid constructs

LAMP1-mGFP and mOrange2-Peroxisomes-2 were gifts from Esteban Dell’Angelica and Michael Davidson (Addgene #34831 and #54596). pPhi-Yellow-mito was obtained from Evrogen (#FP607). LAMP1-PhiYFP was generated by replacing EGFP in LAMP1-mGFP with PhiYFP using BamHI and NotI restriction sites. Clover-mRuby2-FRET-10 and tdTomato-ER-3 were gifts from Michael Davidson (Addgene #58169 and #58097). pAAV-AscI-CAG-H2B-mBaoJin was obtained from WeKwiKgene (#0000270). H2B-Clover was constructed by fusing the H2B sequence from pAAV-AscI-CAG-H2B-mBaoJin with Clover from Clover-mRuby2-FRET-10 into the tdTomato-ER-3 backbone using NheI, Kpn21, and BamHI sites.

#### Live-cell imaging

For live-cell imaging, cells were transiently transfected with LAMP1-PhiYFP, mOrange2-Peroxisomes-2, and H2B-Clover using Lipofectamine 8000 according to the manufacturer’s instructions. Before imaging, cells were incubated with MitoTracker Red CMXRos (500 nM) for 30 min at 37 °C in 5% CO_2_ and with LipidSpot 488 (1:1500) for 5 min at room temperature. Cells were then washed three times with DPBS (5 min each) before imaging.

#### Fixed-cell imaging

For fixed-cell experiments, cells were blocked and permeabilized for 1 h at room temperature in blocking buffer consisting of 3% bovine serum albumin with either 0.5% Triton X-100 or 0.02% saponin in PBS. Samples were incubated overnight at 4 °C with rabbit anti-α-tubulin and mouse anti-Tom20 primary antibodies diluted in blocking buffer. Nuclear nucleic acids were stained with SYBR Gold for 5 min in washing buffer (0.3% BSA and 0.01% Triton X-100 in PBS). After two washes (5 min each), cells were incubated for 1 h at room temperature with goat anti-rabbit IgG–CF514 and goat anti-mouse IgG2a–ATTO542 secondary antibodies. Samples were subsequently washed three times with washing buffer (10 min each) and once with PBS (10 min). After immunostaining, filamentous actin was labeled with YF633-Phalloidin for 20 min in PBS. Finally, LDs were stained with LipidSpot 488 (1:1500) for 5 min, followed by three PBS washes (10 min each) before imaging.

### Morphological library acquisition

A morphological library (*X*_*lmb*_) was constructed to provide structural priors for the adversarial constraint (*L*_*adv*_). Individual subcellular structures were labeled and imaged separately, including nuclei (SYBR Gold), LDs (LipidSpot 488), mitochondria (ATTO542), microtubules (CF514), and actin (YF633), as well as lysosomes and peroxisomes labeled by transient expression of Clover and mOrange2 constructs, respectively. For each organelle class, 70–100 single-channel images were acquired at 512 × 512 pixels. Each image was divided into four non-overlapping 256 × 256 patches, yielding approximately 280–400 structural templates per class. All templates were intensity-normalized before training to improve numerical stability and ensure consistency across labeling modalities.

### Baseline spectral unmixing

To benchmark performance, we compared PhySMI with a classical linear spectral unmixing baseline implemented in an overdetermined regime. Unlike the sparse three-channel input used for PhySMI (*M* = 3), the baseline was provided with the full eight-channel excitation-spectral dataset (*M* = 8) to resolve *N* = 5 fluorophore components.

The multispectral observation *Y* ∈ ℝ^8×*HW*^ was modeled as a linear combination of spectral signatures:

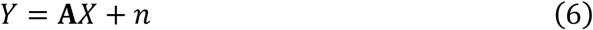

where **A** ∈ ℝ^8×5^ is the experimentally measured excitation spectral signature matrix, *X* denotes the fluorophore abundance maps, and *n* represents measurement noise. To enforce physical plausibility, source abundances were constrained to be non-negative and estimated using non-negative least squares (NNLS)^30^:

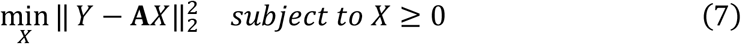

The optimization was solved using an active-set algorithm. By operating with full spectral sampling (*M* > *N*), this baseline represents a conventional linear unmixing reference under favorable acquisition conditions. Comparing PhySMI against this baseline provides a stringent evaluation of its ability to recover latent spectral information and suppress noise from substantially fewer physical measurements.

### Quantitative evaluation metrics

To evaluate reconstruction fidelity and unmixing accuracy, we used four complementary metrics^19, 31^: Peak Signal-to-Noise Ratio (PSNR), Structural Similarity Index (SSIM), Spectral Angle Mapper (SAM), and crosstalk.

PSNR was used to quantify overall reconstruction quality and noise suppression. It was computed from the mean squared error (MSE) between the reference image X and the reconstructed image 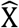:

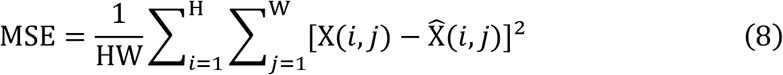

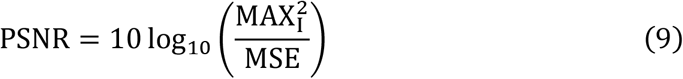

where MAX_I_ is the maximum possible pixel value. Higher PSNR indicates better reconstruction fidelity and stronger suppression of stochastic noise.

SSIM was used to assess preservation of biological morphology by comparing luminance, contrast, and structural similarity between X and the estimate 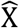. It was computed using a sliding-window implementation:

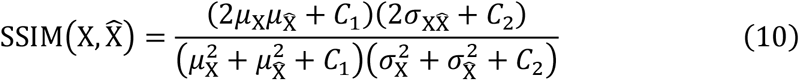

where μ, *σ*, and 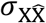 are local means, variances, and covariance, respectively, and *C*_1_, *C*_2_ are constants for numerical stability.

SAM was used to evaluate spectral fidelity by measuring the angular deviation between the estimated spectrum 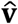 and the reference spectrum **v**:

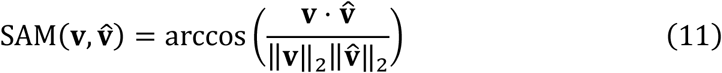

Lower SAM values indicate better spectral agreement and reduced spectral distortion or crosstalk.

To quantify unmixing purity, we calculated crosstalk as the relative leakage of signal into off-target channels. For a target fluorophore *n*, crosstalk into channel *k* (*k* ≠ *n*) within the target mask Ω_*n*_ was defined as:

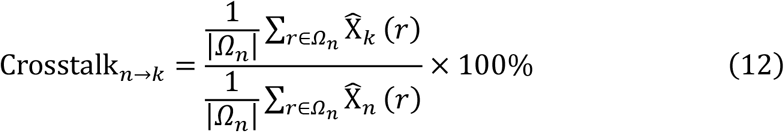

where 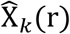 and 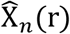 denote the reconstructed intensities of the off-target and target channels at pixel r, respectively. For simulated datasets with spatially isolated targets, this metric directly measures spectral leakage and was visualized as crosstalk matrices under varying noise conditions.

For simulated datasets with spatially isolated targets, any signal reconstructed in an off-target channel within the corresponding target region was considered spectral leakage. Crosstalk was calculated as:

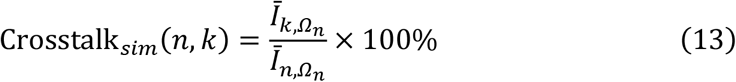

where 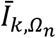 is the mean intensity of channel *k* within the region of the *n*-th target. These values were summarized as crosstalk matrices to assess robustness under varying noise levels.

## Acknowledgments

This work was supported by the National Natural Science Foundation of China (62475032, 62205048), and the Sichuan Science and Technology Program (2026YFHZ0044).

## Author Contributions

K.C. conceived the research. J.X. developed the PhySMI framework and conducted the experiments. All authors contributed to experimental designs, data analysis, and paper writing.

## Code availability

The open-source Python code of PhySMI network is available on GitHub: https://github.com/beiming10/PhySMI.

